# Epigenetic Changes in *Saccharomyces cerevisiae* Alters the Aromatic Profile in Alcoholic Fermentation

**DOI:** 10.1101/2022.08.09.503430

**Authors:** Yanzhuo Kong, Kenneth J. Olejar, Stephen L.W. On, Christopher Winefield, Philip A. Wescombe, Charles S. Brennan, Richard N. Hider, Venkata Chelikani

## Abstract

Epigenetic changes in genomics provide phenotypic modification without DNA sequence alteration. This study shows that benzoic acid, a common food additive and known histone deacetylase inhibitor (HDACi), has an epigenetic effect on *Saccharomyces cerevisiae*. Benzoic acid stimulated formation of epigenetic histone marks H3K4Me2, H3K27Me2, H3K18ac and H3Ser10p in *S. cerevisiae* and altered their phenotypic behavior, resulting in increased production of phenylethyl alcohol and ester compounds during alcoholic fermentation. Our study demonstrates the HDACi activity of certain dietary compounds such as sodium butyrate, curcumin and anacardic acid, suggests the potential use of these dietary compounds in altering *S. cerevisiae* phenotypes without altering host-cell DNA. This study highlights the potential to use common dietary compounds to exploit epigenetic modifications for various fermentation and biotechnology applications as an alternative to genetic modification. These findings indicate that benzoic acid and other food additives may have potential epigenetic effects on human gut microbiota, in which several yeast species are involved.

**Importance:** This manuscript investigates and reports for the first time utilizing microbial epigentics to alter the fermentation process of Pinot noir wines. We have experimentally demonstrated that certain dietary epigenetic compounds possess histone deacetylase (HDAC) inhibiting activity and can alter the wine characteristics by altering yeast gene expression. We have coined the term ‘nutrifermentics’ to represent this newly proposed field of research, which provides insights on the effect of certain dietary compounds on microbial strains and their potential application in fermentation process. This technological approach is a novel way to manipulate microorganisms for innovative food and beverage production with quality attributes.

## 1. Introduction

Epigenetics is the study of phenotypic changes in organisms, which predominantly result from alterations of nucleotides and histones instead of the deoxyribonucleic acid (DNA) sequence; therefore epigenetic modifications are considered a non-GMO approach (1, 2). Among them, DNA methylation and histone acetylation are the most common and well-studied epigenetic modifications, which are the processes of transferring a methyl group to adenine and cytosine, or adding an acetyl group to lysine residues at the N terminus of histone (3). Several dietary bioactive and phytochemicals that naturally occur in fruits and vegetables can act as epigenetic modifiers and potentially can alter the target organism (4). Epigenetic modifiers can be predominantly classified as DNA methyltransferase (DNMT), DNMT inhibitors, histone deacetylase (HDAC), HDAC inhibitors and histone acetyl (HAT) and HAT inhibitors (5, 6).

*S. cerevisiae* is a well-studied model system for epigenetic regulation. Since DNA methylation systems are absent, histone modifications are the primary form of epigenetic regulation, making it a simple system for understanding the relationship between histone modifications and epigenetic states (7, 8). Recently, the fission yeast, *Schizosaccharomyces pombe* was subjected to higher thresholds of caffeine, resulting in epigenetic changes producing transient epimutants with phenotypic plasticity including tolerance to caffeine and cross-resistance to antifungal agents, which was closely related to heterochromatin alterations and heterochromatin-mediated gene silencing (9).

Benzoic acid is a lipophilic weak acid that occurs naturally in many fruits, vegetables, nuts, and even in cultured dairy products as a microbial metabolite (10). Benzoic acid and its derivatives are FDA approved food additives and known histone deacetylase inhibitors (HDACi) that have been shown to stimulate a recently discovered histone mark, lysine benzoylation (11, 12). HDACi compounds play an important role in heterochromatin regulation and gene expression by affecting histone modifications (13).

Here, we investigated the possibility of developing *S. cerevisiae* strains with desirable characteristics for alcoholic fermentation by treating them with the epigenetic modifier, benzoic acid. Benzoic acid was selected due its known capacity to modify histone proteins, its cost-effectiveness and solubility in the aqueous system. We also demonstrate that genes responsible for aroma compounds were upregulated in epimutants compared to the original *S. cerevisiae* strain. The effect of benzoic acid on *S. cerevisiae* H3 histone marks, as benzoic acid is a known HDACi, was also investigated. The results showed that there is several other dietary compounds that could be used to epigenetically alter microbial phenotypes to produce fermented products with desirable characteristics (14, 15).

Wine plays an important role among alcoholic drinks and therefore is representative as a model fermentation system. The wine industry is a competitive industry and developing novel wines is necessary to maintain a competitive advantage in the global market. (16). Compared to some existing approaches, such as grapevine breeding and isolation of wild yeast, epigenetic modification of wine yeast is time and cost effective.

Our study demonstrates the exciting possibility of using dietary epigenetic compounds to develop non-GMO microbial strains with desirable characteristics for fermented products and biotechnology applications.

## 2. Materials and methods

### 2.1 S. cerevisiae starter preparation

A commercial wine yeast *S. cerevisiae* EC-1118 was used as fermentation starters in this study. Three types of starters were involved and applied, including wild type (500 h growth in regular YPD broth, 20 h/sub-culture), epimutant 1 (500 h growth in YPD broth containing 10 mM benzoic acid, 20 h/sub-culture) and epimutant 2 (500 h growth in YPD broth containing 10 mM benzoic acid, followed by 20 h growth in regular YPD broth w/o stress, 20 h/sub-culture). 0.5 mL of cultured broth was transferred for each subculture.

### 2.2 Histone H3 modification multiplex assay

The 21 histone H3 modification patterns of 5 mM benzoic acid treated *S. cerevisiae* compared to untreated wild type strain were measured using EpiQuik™ Histone H3 modification multiplex assay kit (Colorimetric; EpiGentek, NY, USA) following manufacturer’s instructions. Absorbance was measured using FLUOstar Omega microplate reader (BMG LABTECH, Ortenberg, Germany) at 450 nm with a reference wavelength of 655 nm.

### 2.3 RNA purification and gene expression analysis

*S. cerevisiae* under different treatments were harvested after 12 h growth to reach a sample size of 1 × 10^8^ cells for RNA purification. Total RNA from *S. cerevisiae* was isolated using RiboPure™ RNA Purification Kit (Invitrogen, MA, USA), following manufacturer’s instructions. RNA purity was measured by DeNovix DS-11 Spectrophotometer (DeNovix Inc., DE, USA). The RNA expression was measured using nCounter technology (NanoString Technologies, Inc., WA, USA). RNA samples were posted to The University of Auckland, where all the preparation and measurement were completed. Assay was carried out on 12 samples/24 genes (including 5 housekeeping genes), the RNA input amount was 300 ng for each sample. Expression counts were normalized and analyzed using the nSolver 4.0 software (NanoString Technologies, Inc., WA, USA).

### 2.4 DAPI staining

*S. cerevisiae* strains were cultured overnight to an OD_600_ = 1.0 ± 0.2, followed by being treated with 2 volumes of 100% Ethanol for 45 min at room temperature. The mixture was centrifuged at 2500 rpm for 1 min, 1 mL 1 × PBS was used to wash the cells, followed by another centrifugation at 2500 rpm for 1 min. The pellet was resuspended in 200 µL of 1 × PBS/1:2000 dilution DAPI mixture, and was observed under Nikon Eclipse 50i fluorescence microscope (Nikon, Tokyo, Japan) after 45 min.

### 2.5 Yeast morphology

*S. cerevisiae* starters were transferred from YPD broth onto corresponding YPD agar plates. Cultured media were serially diluted to OD_600_ = 0.1, 5 µL strain solution was spotted onto corresponding YPD agar plates after an additional 10 times dilution being applied, 1 × PBS was used for dilution. The growth temperature was set at 32 °C.

### 2.6 GC-MS and chemical analysis of wine samples

The alcohol and ester aroma compounds analysis was conducted using headspace-solid phase microextraction (HS-SPME) and Shimadzu QP-2010 GC-MS (Shimadzu, Kyoto, Japan). The methodology was adopted from previous published articles, with slight modification regarding the diluent and sample matrix used with the standards (17, 18). Detailly, 0.9 mL of sample was pipetted into a 20 mL amber SPME vial and diluted with 8.06 mL of 5 g/L tartaric acid buffer (pH 3.5), 40 µL of composite internal standard was added followed by 4.5 g of sodium chloride before the vial was immediately capped.

For the preparation of the highest standard of the calibration curve, the composite standard was diluted in 136 mL sample matrix which was rotary evaporated at 36 °C for 40 min to remove volatile background, and reconstituted with 14.2% Ethanol as well as 40 µL of 5 M sodium hydroxide which returned the pH back to 3.15. It was then serially diluted in the provided matrix to ensure each vial had a maximum volume of 0.9 mL of matrix present. Each vial was then diluted further with 8.06 mL of tartaric acid buffer as in the samples with 40 µL of composite internal standard being added, followed by 4.5g of sodium chloride before the vials were immediately capped.

Ethanol content was analyzed by GC-FID, which was carried out on a Shimadzu GC-2010 gas chromatograph-flame ionization detector equipped with an AOC-20i autoinjector and AOC-20s autosampler. The chromatography was performed using an 19091N-133 HP-Innowax GC column (PoAgilent Technologies, CA, USA). Residual sugars including glucose and fructose were measured using Vintessential enzymatic test kit (Vintessential Laboratories – Tasmania, TAS, Australia), and glycerol content was measured using Megazyme glycerol assay kit (Megazyme, Wicklow, Ireland).

### 2.7 HDAC inhibition assay

The HDAC inhibition capacity of candidate epigenetic modifiers was measured by a fluorometric HDAC assay kit (Active Motif, Inc., CA, USA), following the manufacturer’s instructions with slight modification to suit the objectives of this research. HeLa nuclear extract was used as the HDAC source, with an input volume of 5 µL. Candidate epigenetic modifiers/HDAC inhibitors, including the positive control Trichostatin A (TSA), were added at the volume of 10 µL. The volume of HDAC assay buffer was adjusted to reach a total volume of 50 µL in each well. Fluorescence was measured using FLUOstar Omega microplate reader (BMG LABTECH, Ortenberg, Germany) with excitation wavelength at 360 nm and emission wavelength at 460 nm.

### 2.8 Statistical analysis

Results were gathered from three independent biological replicates unless otherwise stated. Data were analyzed using analysis of variance (ANOVA) with a generalized linear model, followed by *post-hoc* Tukey’s mean comparison test, using Minitab 20 (Minitab, LLC, PA, USA). PCA and AHC were analyzed using XLSTAT Statistical Software 2016 (Addinsoft, Paris, France). A confidence level of 95% was applied to the statistical analysis and data are presented as mean ± SD.

## 3. Results and discussion

### 3.1 Project scope: the practice of altering fermentation by epigenetics

The schematic diagram put forward to fit the entire scope of the project is shown in Figure 1 and demonstrates the proposed innovation to food fermentation by impacting gene expression levels of microbial starters using diet-derived epigenetic modifiers. There are a range of diet-derived epigenetic modifiers including bioactive compounds and phytochemicals, which are of health benefits to humans. For example, diet derived short-chain fatty acids are a group of HDAC inhibitors which are known to play a key role in epithelial hemostasis and repair process (14). The research investigating the effect of food and food components on gene expression, and its role involved in the interaction between host/microbes and the nutritional environment is a well-established research field called “nutrigenomics” (Figure 1A). In this study, we have shown that we can use these dietary epigenetic compounds such as dietary HDACi to alter the microbial phenotypes used in the fermentation process. We coined the word “nutrifermentics” (Figure 1B) to represent this new field of research. These dietary HDACi could also provide health benefits to consumers, in addition to improving starter microbial cultures in the fermentation process.

**Figure 1.**
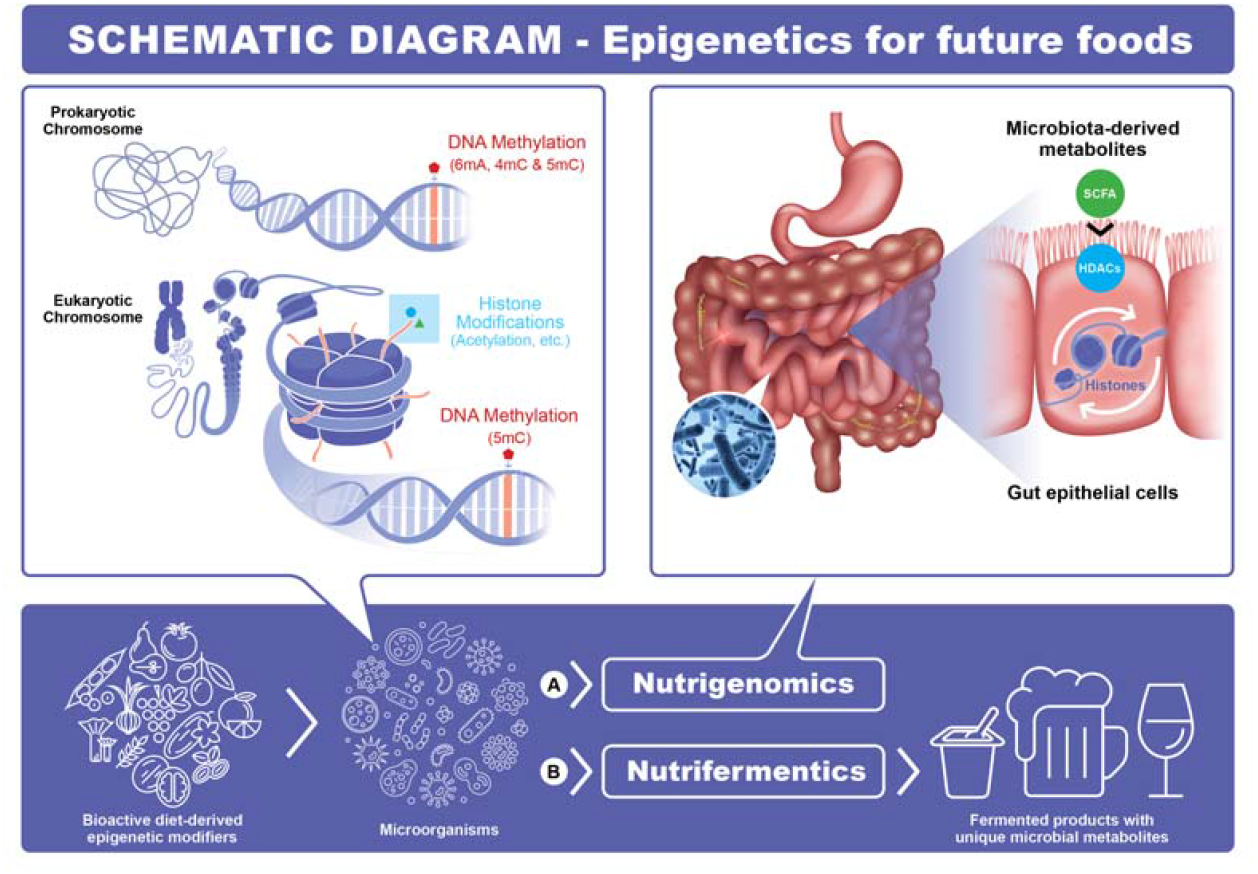
Schematic diagram: An innovation to food fermentation by impacting gene expression levels of microbial starters using diet-derived epigenetic modifiers. **A) Nutrigenomics:** A well-established field of research investigating the effect of food and food components on gene expression, and its role involved in the interaction between host/microbes and the nutritional environment; **B) Nutrifermentics:** A new research direction firstly proposed in this study, which provides insights regarding the effect of food and food components on microbial starters and its potential application in fermentation. **Abbreviations: SCFA:** short-chain fatty acids; **HDACs:** histone deacetylases.

### 3.2 Influence of benzoic acid on *S. cerevisiae* histone H3

Benzoic acid and its derivatives are known HDACi (11) and recent study revealed that sodium benzoate can stimulate a new histone mark, lysine benzoylation with significant physiological relevance (12). Figure 2A shows the percentage of relative changes in 21 distinct histone H3 modification patterns in *S. cerevisia,e* which was treated with 5 mM benzoic acid, in comparison with untreated wild type strain. Specific antibodies, including 15 for methylation, 4 for acetylation and 2 for phosphorylation were utilized to measure the 21 patterns. Most modification patterns between treated and untreated strains were around 100% when taking the variation into consideration. However, both stimulation and inhibition in histone marks were seen with exposure to 5 mM benzoic acid. H3K4me2, H3K9me3, H3K27me2, H3K9ac, H3K18ac and H3ser10p were stimulated more than four-fold in treated strains, whereas few methylation patterns including H3K4me3, H3K9me2 and H3K27me3 were about half compared to the untreated strain. Histone modifications are directly relevant to gene expression levels in the organism

**Figure 2.**
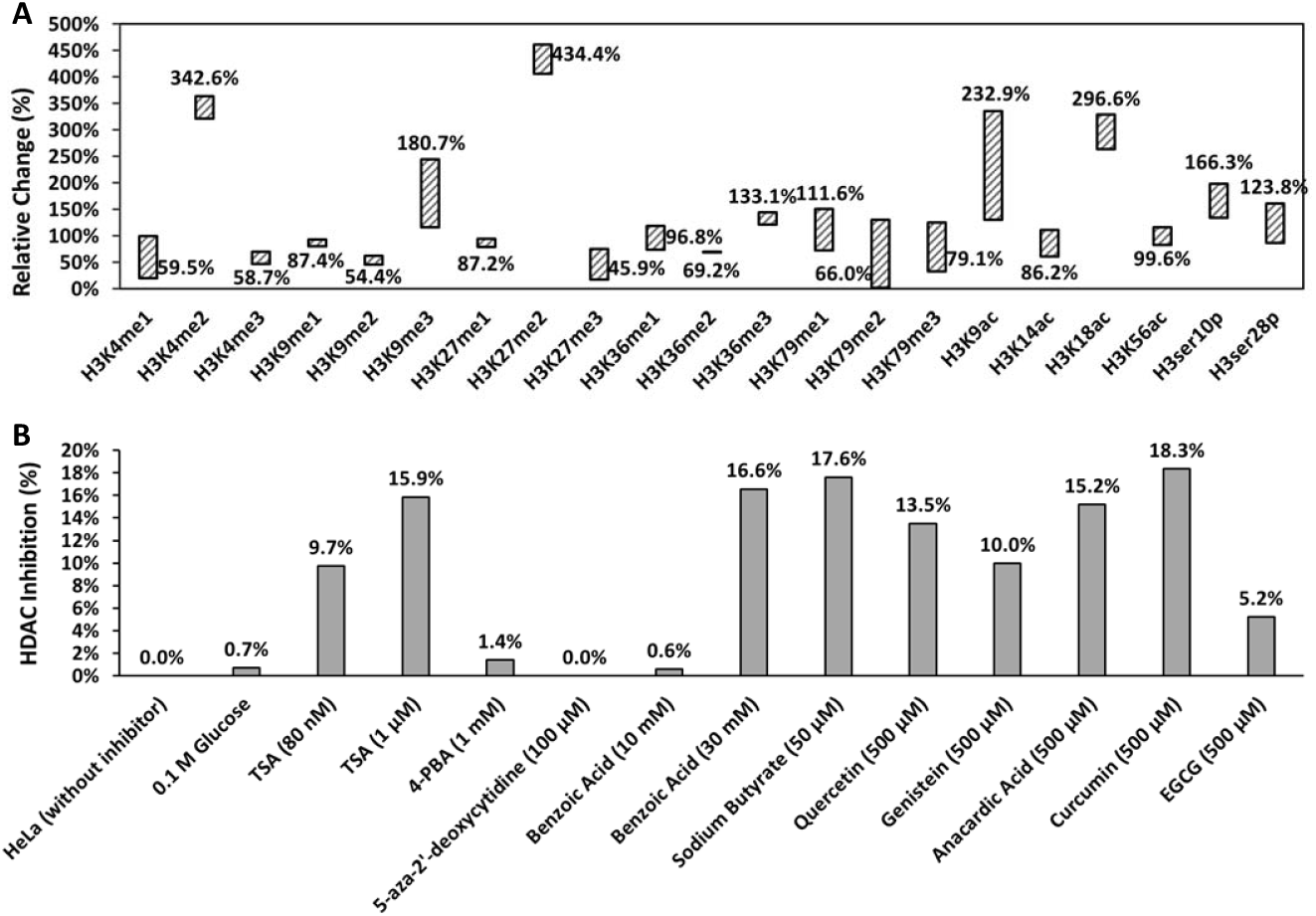
**A**. Relative changes of 21 histone H3 modification patterns between *S. cerevisiae* wild type and strains treated with 5 mM benzoic acid. **B**. Histone deacetylase (HDAC) inhibition capacity of candidate epigenetic modifiers at different concentrations, in comparison with HeLa cells without inhibitors. **Abbreviations: TSA:** trichostatin A; **4-PBA:** 4-phenylbutyric acid; **EGCG:** epigallocatechin gallate.

### 3.3 Gene expression analysis by NanoString

NanoString transcription analysis revealed the expression of 24 genes including five housekeeping genes. Figure 3 shows the gene expression levels of *S. cerevisiae* under different treatments, TSA treatments including first-time exposure, 500 h treatment (20 h/sub-culture) and 1 generation w/o treatment after 500 h exposure were included as the HDACi controls, which were in accordance with benzoic acid treatments (first-time exposure to 5 mM benzoic acid, 500 h treatment (20 h/sub-culture) at 10mM benzoic acid /epimutant 1 and 1 generation w/o treatment after 500h exposure/epimutant 2). The untreated wild type strain was included as a negative control along with 0.9% sodium chloride and with a dietary polyphenol epigallocatechin gallate (EGCG) that has been reported to inhibit DNMTs (4).

**Figure 3.**
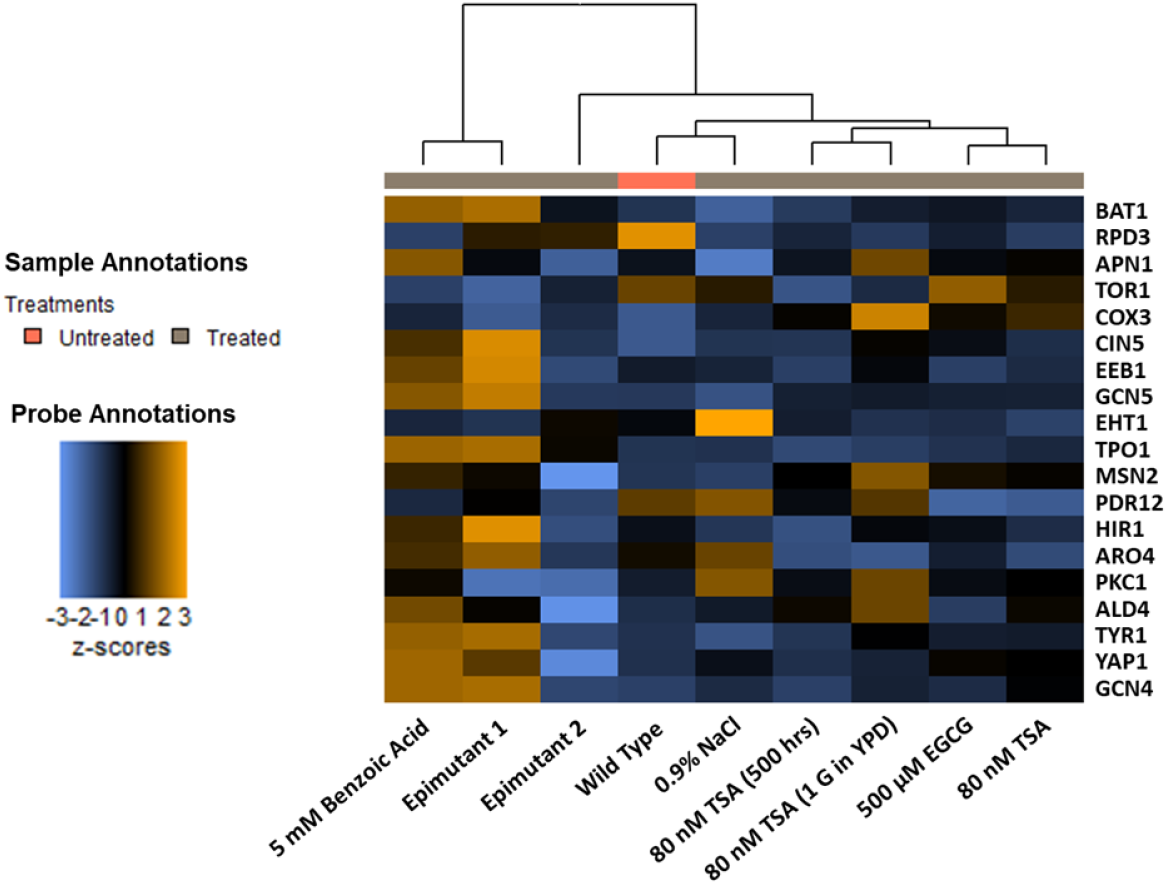
Heat map showing unsupervised hierarchical clustering of 9 S. cerevisiae samples under different treatments, based on their expression levels of 19 selected genes. **Abbreviations: TSA:** trichostatin A; **EGCG:** epigallocatechin gallate.

Results are presented as a heat map graph after Z-score transformation, ranging from -3 to 3, blue (downregulation) to orange (upregulation), Figure 3. As shown in Figure 3, RNA samples with different treatments were clustered after data normalization, in which benzoic acid supplementation became a distinct influencing factor.

Genes responsible for overproducing phenylethyl alcohol (ARO4 and TYR1) (19), fusel alcohol and ester synthesis (EEB1) (20), biosynthesis of higher alcohols (BAT1) (21) were all upregulated. There was no significant change in the expression levels of another ester synthase gene (EHT1) (20) between samples. Several genes responsible for stress tolerance and cell cycle were analyzed finding the histone deacetylase gene (RPD3) (22) expression was downregulated while the histone acetyltransferase gene (GCN5) (23) was upregulated in both 5mM benzoic acid-treated strain and epimutant 1, clearly supporting the role of benzoic acid as an HDACi. Kurat et al. (23), previously reported that upregulation of the GCN5 expression led to histone acetylation and global transcriptional activation. The variation observed could potentially indicate alternative acetylation mechanisms in *S. cerevisiae* resulted from different concentrations and exposure time of certain epigenetic compounds such as benzoic acid. With respect to the clusters, RNA from *S. cerevisiae* epimutant 1 (500 h growth in YPD broth containing 10 mM benzoic acid, 20 h/sub-culture) showed quite similar expression patterns to 5 mM benzoic acid treatment (first time exposure). Moreover, the epimutant 2 (500 h growth in YPD broth containing 10 mM benzoic acid, followed by 20 h growth in regular YPD broth w/o stress, 20 h/sub-culture) exhibited significantly different RNA expression patterns compared with the benzoic acid treatment group. The epimutant 2 tended to be more relevant to wild type and other *S. cerevisiae* treatments. This observation suggests that the alteration of gene expression caused by dietary epigenetic compounds, which is revealed by direct counts of RNA transcripts, is transient and tends to ease out once the stress inducer is eliminated from the environment.

### 3.4 Influence of benzoic acid on *S. cerevisiae* nucleus

As shown in Figure 4A, DAPI (4’, 6-diamidino-2-phenylindole) staining was applied to wild type and epimutant 1 to visualize their nuclei in terms of any size changes (expansion) that may have resulted from benzoic acid treatment. As DAPI stoichiometrically binds to DNA, which enables the detection and comparison of DNA content variation by fluorescence microscopy (24). The corrected total cell fluorescence (CTCF) was calculated based on the integrated density in the nucleus region. The mean comparison results indicate that there is a significant difference between two samples (*p* < 0.05), which suggests an expansion to the nucleus region in *S. cerevisiae* when they are exposed to 10 mM benzoic acid. A. D. Walters et al. (25) suggested that the expansion of nuclear envelope in budding yeast is independent from cell growth, but potentially related to nucleoplasmic factors, such as one or more nucleoplasmic proteins that are synthesized or imported into the nucleus. As HDAC and HDACi have been well researched being involved in multiple cell processes, such as cytokinesis and apoptosis (26, 27). Therefore benzoic acid induced, HDACi related modifications could have occurred in nucleoplasm, resulting in expanded nucleus or relaxed genome state, which led to more fluorescence in this study.

**Figure 4.**
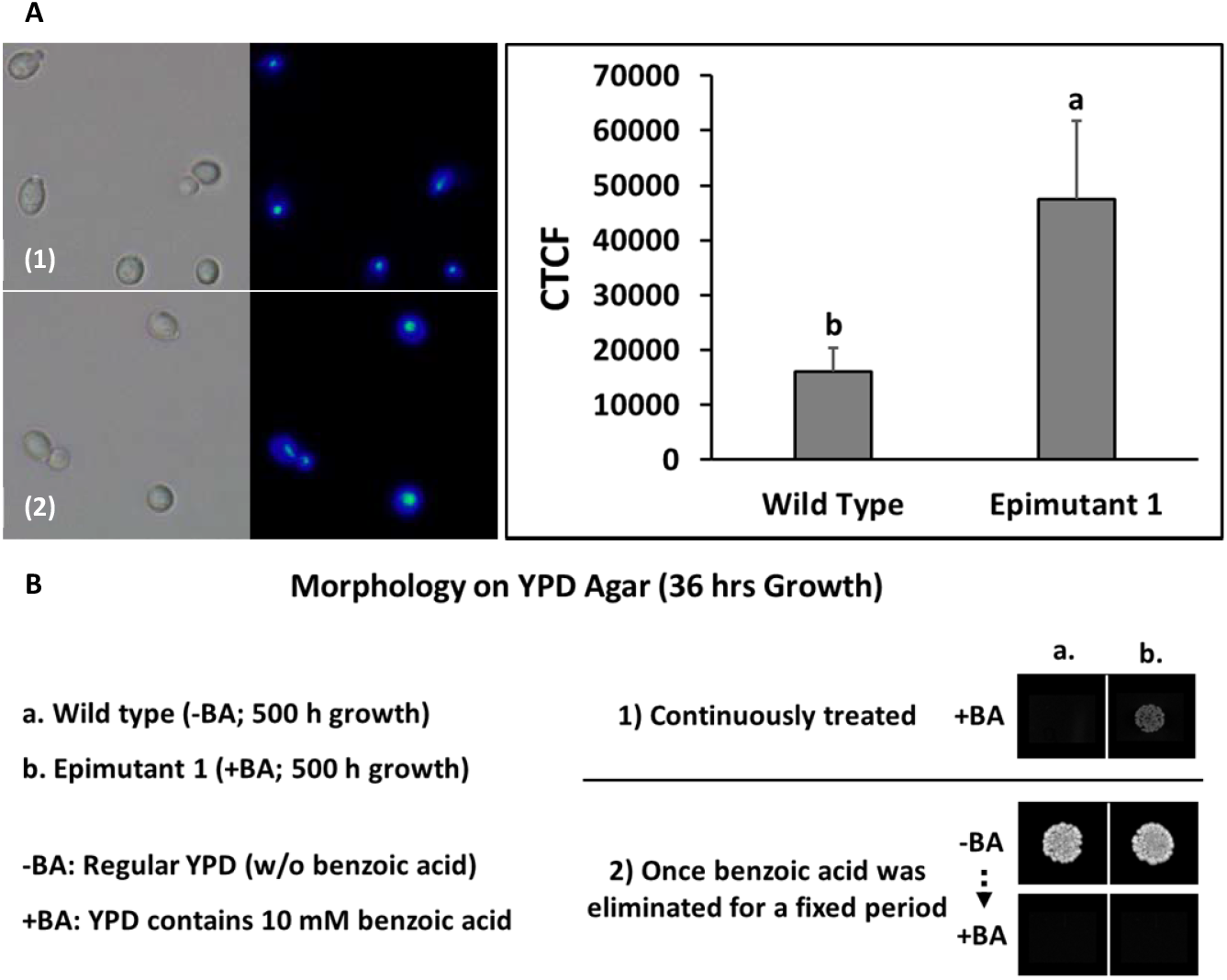
**A**. DAPI staining (bright-field/fluorescent, at 100X) and corrected total cell fluorescence levels of benzoic acid-treated *S. cerevisiae* in comparison with wild type. **B**. Phenotypic plasticity of benzoic acid-treated *S. cerevisiae* epimutant. **(1):** Wild Type; **(2):** Epimutant 1; **CTCF:** corrected total cell fluorescence; **a-b:** different letters indicate significant difference based on Tukey pairwise mean comparison results (*p* < 0.05).

### 3.5 Yeast morphology in relation to epigenetic alteration

The cellular morphology of benzoic acid-treated *S. cerevisiae* was recorded to depict phenotypic differences compared to the wild type (Figure 4B). Epimutant 1 was more tolerant to benzoic acid treatment and showed visible alteration in colony morphology. Observed tolerance and adaptation to benzoic acid faded soon after the stress was eliminated from the environment, demonstrating the transient nature of the treatment and possibly epigenetic change. The observation is supported by S. Torres-Garcia et al. (9); phenotypic plasticity can be promoted by epigenetic processes that let the wild type cells adapt to certain unfavorable environments without altering genetic information, although these alterations are generally unstable and will be gradually lost without the stress. This observation is in line with NanoString assay, where *S. cerevisiae* epimutant 1 showed very similar expression patterns to 5 mM benzoic acid-treated strain (first time exposure). However, epimutant 2 exhibited significantly different gene expression patterns when compared within the benzoic acid treatment group. Epimutant 2 gene expression patterns were more similar to wild type than other treatments. This suggests that the alteration of gene expression caused by dietary epigenetic compounds is transient and tends to fade once the compound is eliminated from the environment. Overall, the robustness of *S. cerevisiae* epimutants and their adaption to stressed environment were improved by continuously treating the strain with the threshold levels of benzoic acid. However, epigenetic plasticity could be an issue in retaining the robust characteristics for future generations once the epigenetic modifiers is removed from the environment. However, commercial yeast starter culture producer could potentially prefer single/double use strains similar to commercial seed companies.

### 3.6 Wine characteristics changes due to epigenetic alteration

To test the impact of benzoic acid-stimulated epigenetic changes on fermentation characteristics of *S. cerevisiae*, treated cultures were used to ferment wine samples. Wines were fermented using three *S. cerevisiae* starters, including wild type, epimutant 1 (500 h growth in YPD broth containing 10 mM benzoic acid, 20 h/sub-culture) and epimutant 2 (epimutant 1 followed by 20 h growth in regular YPD broth without stress). Principal component analysis (PCA) of aromatic attributes of wine samples and agglomerative hierarchical clustering (AHC) were utilized for classification of fermented wine samples (Figure 5A). In addition, GC-MS analysis was carried out on wine samples, distinct results are shown in Figure 5B, with full analysis of 18 compounds listed in Table 1. The three starters resulted in three wine categories, each with distinct aromatic profiles (Figure 5A & 5B). The positive correlation between epimutant 1 and *cis*-3-hexen-1-ol may indicate a kiwifruit and leaf-like aroma is potentially associated with wine produced by epimutant 1 (28). Since *cis*-3-hexen-1-ol is an important aroma compound in many white wines, it might confer a complex aromatic profile on the altered wine, by adding partial aromatic features of white wine. As shown in Figure 5B, the content of five ester and higher alcohol compounds is listed as potential indication of wine aroma alterations resulting from epimutation of the starters. Wine fermented by epimutant 1 possessed significantly increased phenylethyl alcohol (rose scent), ethyl lactate (butter aroma), *cis*-3-hexen-1-ol (leaf alcohol confers grassy-green odor) and ethyl pentanoate (fruity aroma), whereas the content of ethyl octanoate was reduced (soapy, floral aroma) (*p* < 0.05) (29). This GC-MS analysis clearly supports the gene expression observed using Nanostring assay. This is a significant advancement in the fermentation field considering the treatments lead to over expression of genes associated with favorable aromas such as Phenyl ethyl alcohol and overproduction of these compounds are confirmed by GC-MS analysis. In addition to aroma alterations, major chemical composition changes due to epigenetic alterations were also investigated, including residual sugars (glucose and fructose), glycerol and ethanol (Supplementary Dataset 1). Generally, epimutant 1 tended to increase the fructose content in wine (*p* < 0.05), whereas not significantly affect the content of other chemicals analyzed (*p* > 0.05). The study detected a few more distinct aromatic compounds by qualitative GC-MS in these wine samples and is listed in Supplementary Dataset 2.

**Table 1.**
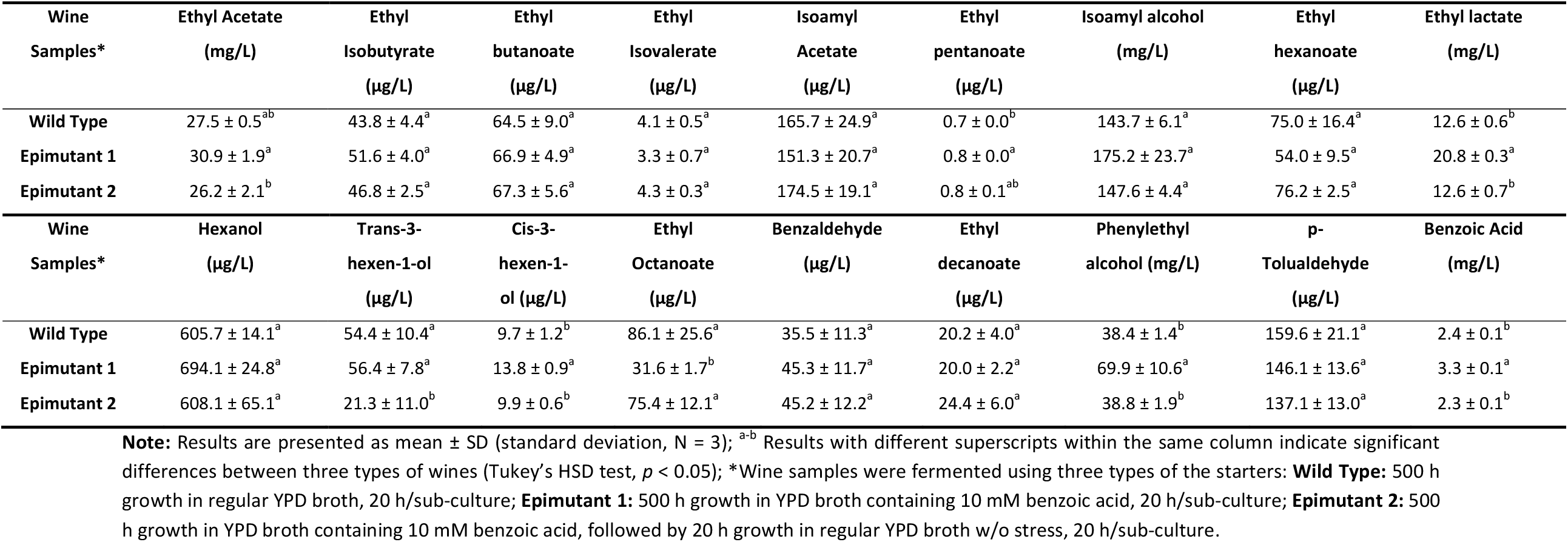
Analysis of alcohol and ester aroma compounds detected in Pinot Noir wine.

| Wine Samples* | Ethyl Acetate (mg/L) | Ethyl Isobutyrate (µg/L) | Ethyl butanoate (µg/L) | Ethyl Isovalerate (µg/L) | Isoamyl Acetate (µg/L) | Ethyl pentanoate (µg/L) | Isoamyl alcohol (mg/L) | Ethyl hexanoate (µg/L) | Ethyl lactate (mg/L) |
| --- | --- | --- | --- | --- | --- | --- | --- | --- | --- |
| Wild Type | 27.5 ± 0.5<sup>ab</sup> | 43.8 ± 4.4<sup>a</sup> | 64.5 ± 9.0<sup>a</sup> | 4.1 ± 0.5<sup>a</sup> | 165.7 ± 24.9<sup>a</sup> | 0.7 ± 0.0<sup>b</sup> | 143.7 ± 6.1<sup>a</sup> | 75.0 ± 16.4<sup>a</sup> | 12.6 ± 0.6<sup>b</sup> |
| Epimutant 1 | 30.9 ± 1.9<sup>a</sup> | 51.6 ± 4.0<sup>a</sup> | 66.9 ± 4.9<sup>a</sup> | 3.3 ± 0.7<sup>a</sup> | 151.3 ± 20.7<sup>a</sup> | 0.8 ± 0.0<sup>a</sup> | 175.2 ± 23.7<sup>a</sup> | 54.0 ± 9.5<sup>a</sup> | 20.8 ± 0.3<sup>a</sup> |
| Epimutant 2 | 26.2 ± 2.1<sup>b</sup> | 46.8 ± 2.5<sup>a</sup> | 67.3 ± 5.6<sup>a</sup> | 4.3 ± 0.3<sup>a</sup> | 174.5 ± 19.1<sup>a</sup> | 0.8 ± 0.1<sup>ab</sup> | 147.6 ± 4.4<sup>a</sup> | 76.2 ± 2.5<sup>a</sup> | 12.6 ± 0.7<sup>b</sup> |

| Wine Samples* | Hexanol (µg/L) | Trans-3-hexen-1-ol (µg/L) | Cis-3-hexen-1-ol (µg/L) | Ethyl Octanoate (µg/L) | Benzaldehyde (µg/L) | Ethyl decanoate (µg/L) | Phenylethyl alcohol (mg/L) | p-Tolualdehyde (µg/L) | Benzoic Acid (mg/L) |
| --- | --- | --- | --- | --- | --- | --- | --- | --- | --- |
| Wild Type | 605.7 ± 14.1<sup>a</sup> | 54.4 ± 10.4<sup>a</sup> | 9.7 ± 1.2<sup>b</sup> | 86.1 ± 25.6<sup>a</sup> | 35.5 ± 11.3<sup>a</sup> | 20.2 ± 4.0<sup>a</sup> | 38.4 ± 1.4<sup>b</sup> | 159.6 ± 21.1<sup>a</sup> | 2.4 ± 0.1<sup>b</sup> |
| Epimutant 1 | 694.1 ± 24.8<sup>a</sup> | 56.4 ± 7.8<sup>a</sup> | 13.8 ± 0.9<sup>a</sup> | 31.6 ± 1.7<sup>b</sup> | 45.3 ± 11.7<sup>a</sup> | 20.0 ± 2.2<sup>a</sup> | 69.9 ± 10.6<sup>a</sup> | 146.1 ± 13.6<sup>a</sup> | 3.3 ± 0.1<sup>a</sup> |
| Epimutant 2 | 608.1 ± 65.1<sup>a</sup> | 21.3 ± 11.0<sup>b</sup> | 9.9 ± 0.6<sup>b</sup> | 75.4 ± 12.1<sup>a</sup> | 45.2 ± 12.2<sup>a</sup> | 24.4 ± 6.0<sup>a</sup> | 38.8 ± 1.9<sup>b</sup> | 137.1 ± 13.0<sup>a</sup> | 2.3 ± 0.1<sup>b</sup> |

**Note:** Results are presented as mean ± SD (standard deviation, N = 3); <sup>a-b</sup> Results with different superscripts within the same column indicate significant differences between three types of wines (Tukey's HSD test, $p < 0.05$ ); \*Wine samples were fermented using three types of the starters: **Wild Type:** 500 h growth in regular YPD broth, 20 h/sub-culture; **Epimutant 1:** 500 h growth in YPD broth containing 10 mM benzoic acid, 20 h/sub-culture; **Epimutant 2:** 500 h growth in YPD broth containing 10 mM benzoic acid, followed by 20 h growth in regular YPD broth w/o stress, 20 h/sub-culture.

**Figure 5.**
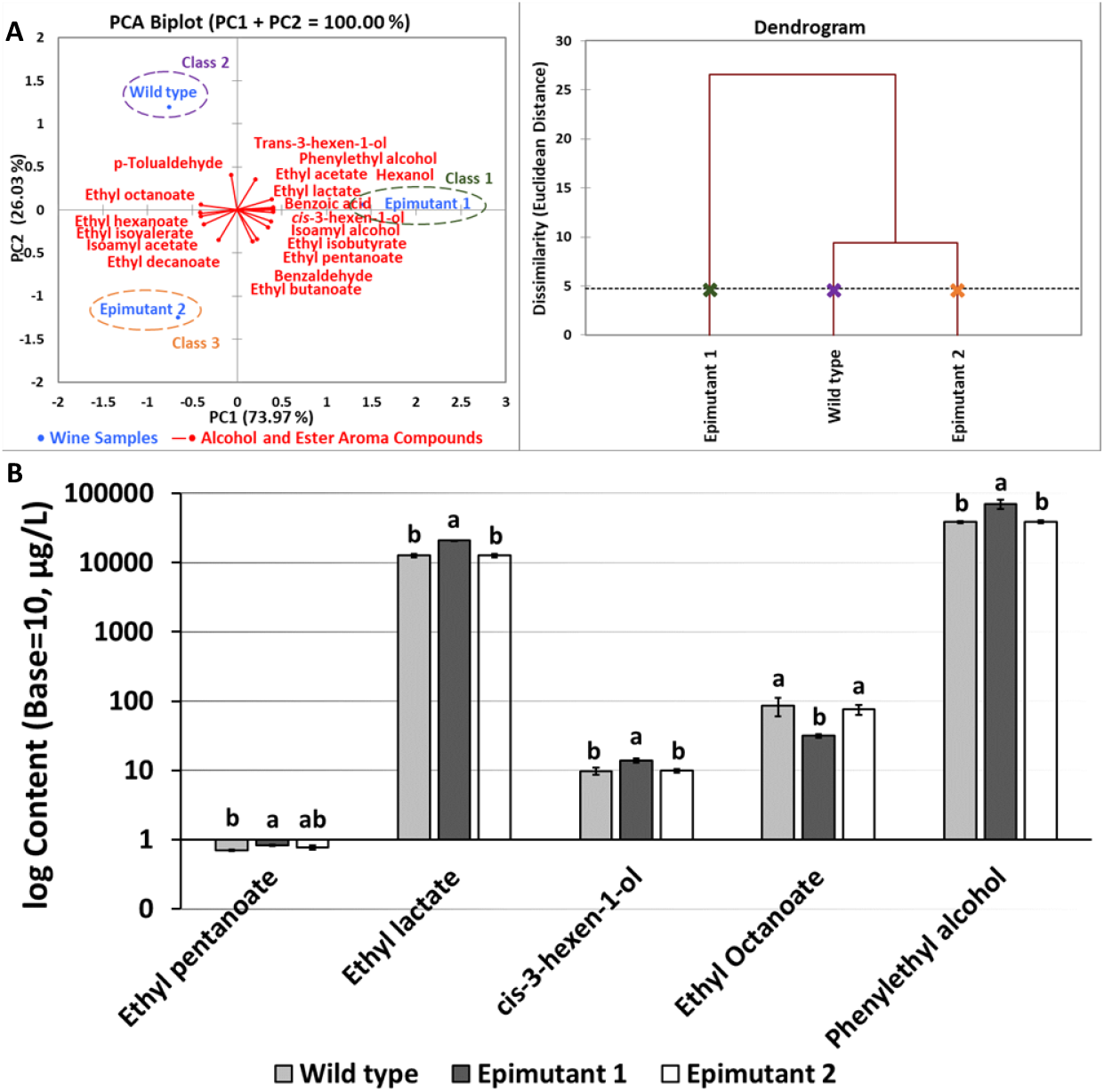
**A. Left:** Principal component analysis (PCA) bi-plot illustrating the relationship between wine samples fermented under different conditions and the variance of alcohol and ester aroma compounds; **Right:** Wine samples grouped using agglomerative hierarchical clustering (HCA) according to dissimilarity levels based on GC-MS analysis. **B**. The content of ester and higher alcohol compounds potentially contributing to distinct aromatic profiles of wine samples fermented by epimutated *S. cerevisiae*. **a-b:** different letters indicate significant difference based on Tukey pairwise mean comparison results (*p* < 0.05).

### 3.7 HDAC inhibition capacity

As in previous NanoString assay, significantly different patterns of transcribed genes by other treatments compared to benzoic acid group, suggesting that HDACi and compounds capable of modifying epigenetic states have different effects on histone proteins and gene expression patterns, potentially indicating wide range of application for these HDACi (Figure 3). This led us to consider the HDACi activity of dietary epigenetic compounds. To investigate this we applied a well-established HDAC assay using HeLa cell lines (14, 15). The dietary compounds tested were sodium butyrate, quercetin, genistein, anacardic acid, curcumin and EGCG (Figure 2B). Untreated HeLa nuclear extracts were used as the negative control. 5-Aza-2’-deoxycytidine, which is a well-recognised DNMT inhibitor but not HDAC inhibitor (30), and glucose were also included as negative controls for assay calibration. A well-known HDACi, namely TSA, was used as a generic inhibitor of histone acetylation to test the impact on gene transcription (31). Relevant half-maximal inhibitory concentrations (IC_50_) were referred for determining the testing concentrations of candidate chemicals, except for benzoic acid since an IC_50_ was not determined. In general, most of the tested dietary compounds exhibited equivalent or better HDACi capacity compared to TSA, the positive control, suggesting their potential application in the food industry, particularly the food fermentation field.

## 4. Conclusions

This study showed the potential applications of dietary epigenetic compounds in food research. As a proof of concept, it has been shown that epigenetic changes in yeast *S. cerevisiae* can be induced using dietary compounds resulting in different aromatic profiles in alcoholic fermentation. This opens the exciting possibility of using a non-GMO approach to obtain microbial strains with desirable characteristics for fermented food products. Interestingly, it was observed that the downregulation of H3K27me3 histone mark and upregulation of GCN4 gene, which are associated with life span extension in *C*.*elgans* and *S. cerevisiae*, respectively (32-34). Understanding the role of these dietary epigenetic compounds on cell ageing is potentially an interesting future research.

## Author Contributions

**Yanzhuo Kong:** Investigation, Validation, Formal Analysis, Writing – Original Draft & Writing – Review & Editing; **Kenneth J. Olejar:** Investigation, Validation & Writing – Review & Editing; **Stephen L.W. On:** Validation & Writing – Review & Editing; **Christopher Winefield:** Validation & Writing – Review & Editing; **Philip A. Wescombe:** Validation & Writing – Review & Editing; **Charles S. Brennan:** Writing – Review & Editing; **Richard N. Hider:** Investigation & Validation; **Venkata Chelikani:** Conceptualization, Writing – Review & Editing, Resources, Supervision & Funding Acquisition.

## Acknowledgments

The work described is in relation to Australian Provisional Patent Number: 2022901674. We thank Shibani Suresh for her contributions to the initial part of the research, Jason Breitmeyer for his technical and experimental support towards GC-MS analysis, and Lincoln University Marketing and Customer Engagement Team for designing the schematic diagram.

